# A Comprehensive Tool Set for Inducible, Cell Type-Specific Gene Expression in Arabidopsis

**DOI:** 10.1101/242347

**Authors:** Ann-Kathrin Schürholz, Vadir Lopez-Salmeron, Zhenni Li, Joachim Forner, Christian Wenzl, Christophe Gaillochet, Sebastian Augustin, Amaya Vilches Barro, Michael Fuchs, Michael Gebert, Joop E.M. Vermeer, Jan U. Lohmann, Thomas Greb, Sebastian Wolf

## Abstract

Understanding the context-specific role of gene function is a key objective of modern biology. To this end, we generated a resource for inducible cell-type specific trans-activation based on the well-established combination of the chimeric GR-LhG4 transcription factor and the synthetic *pOp* promoter. Harnessing the flexibility of the GreenGate cloning system, we produced a comprehensive set of GR-LhG4 driver lines targeting most tissues in the *Arabidopsis* shoot and root with a strong focus on the indeterminate meristems. We show that, when combined with effectors under control of the *pOp* promoter, tight temporal and spatial control of gene expression is achieved. In particular, inducible expression in F1 plants obtained from crosses of driver and effector lines allows rapid assessment of the cell type-specific impact of an effector with high temporal resolution. Thus, our comprehensive and flexible toolbox is suited to overcome the limitations of ubiquitous genetic approaches, the outputs of which are often difficult to interpret due to widespread existence of compensatory mechanisms and the integration of diverging effects in different cell types.

One sentence summary: A set of lines enabling spatio-temporal control of gene expression in Arabidopsis.

## Introduction

The key to the evolutionary success of multicellularity, which arose independently in plants and animals, is the division of labour between highly specialized cell types. This requires the robust specification of cell fate through epigenetic and transcriptional programming, despite the identical genetic makeup of each cell. In plants, cell fate acquisition is largely based on positional information, which depends on cell-to-cell communication and medium to long distance morphogenetic signals that cooperate in organ patterning (Efroni, 2017). Conversely, individual genes, pathways, and metabolites can have diverse or even opposing roles depending on the tissue context. A prominent example for context-dependency of a fundamental patterning process is given by the interplay of the auxin and cytokinin phytohormones (Furuta et al., 2014; Greb and Lohmann, 2016; Truskina and Vernoux, 2017). In the shoot apical meristem, harbouring the stem cell niche ultimately responsible for most above-ground plant organs, cytokinin signalling is associated with maintaining a pluripotent, “undifferentiated” state, whereas auxin signalling promotes differentiation. In marked contrast, auxin is required for stem cell maintenance in the root apical meristem (RAM) (Pacifici et al., 2015; Weijers and Wagner, 2016). Therefore, the global effects of genetic lesions or of knockins can dilute and mask specific functions and are often difficult to interpret.

Routinely, stable genetic gain- and loss-of-function mutants remain the main pillar of the reductionist approach to biology and the phenotypes of such mutants are assessed to deduce a function of the mutated locus in wildtype. However, in addition to the context-dependency of many gene products, the manifest phenotypes of mutants or transgenic lines are derived from an unknown combination of primary and secondary effects caused by the genetic alteration. Thus, mutant organisms can undergo life-long adaptation and compensation processes usually unbeknownst to the experimenter, impeding the interpretation of their phenotype. In addition, transgenic and mutational approaches can interfere with plant vitality, precluding an in-depth analysis.

Many of these problems can be overcome by inducible, cell type-specific expression mediated by two-component transcription activation systems (Moore et al., 2006). An expression cassette is constructed using a heterologous or synthetic promoter and is hence silent unless a cognate transcription factor is present. An efficient approach is to generate driver lines that express the transcription factor in a spatially and temporally controlled manner on the one hand, and a responder line carrying the effector construct on the other hand. After crossing of the two lines, expression can be induced and the phenotypic consequences of the effector can be studied. In the abstract, these expression systems are highly valuable because they ideally enable cell-type specific or stage-specific complementation or knock-down, facilitate time-resolved monitoring of the response to a given cue, can overcome lethality of constitutive expression, and allow to study cell autonomous and non-cell autonomous effects with high temporal and spatial resolution. However, the considerable effort and time requirements for DNA cloning and the generation of stable transgenic plants are a major bottleneck curtailing their use to date. For the same reason and because distinct tissue-specific promoters were not always available in the past, attention is usually given to one tissue or cell type of interest at a time and unbiased approaches targeting a larger spectrum of individual tissues are hardly followed.

Here, we report on the generation of a comprehensive set of *Arabidopsis* driver lines suited for tissue specific trans-activation of an effector cassette in a wide range of cell types and with the possibility to monitor gene activation in space and time by a fluorescent promoter reporter. To ensure rapid, stable induction with minimal adverse effects on plant growth caused by the inducer, our system takes advantage of the widely used LhG4/pOp system (Moore et al., 1998; Craft et al., 2005; Samalova et al., 2005) combined with the ligand binding domain of the rat glucocorticoid receptor (GR) (Picard, 1993) (Craft et al., 2005). LhG4 is a chimeric transcription factor consisting of a mutant version of the *Escherichia coli lac* repressor with high DNA binding affinity (Lehming et al., 1987) and the transcription activation domain of yeast Gal4p (Moore et al., 1998). N-terminal fusion with the GR ligand binding-domain renders the transcription factor inactive in the cytosol through sequestration by HSP90 in the absence of the inducer. Nuclear import after treatment with the synthetic glucocorticoid dexamethasone (Picard, 1993) results in transcriptional activation of expression cassettes that are under control of the synthetic Op 5’ regulatory region consisting of a Cauliflower Mosaic Virus (CaMV) *35S* minimal promoter and two upstream *lac* operators (Moore et al., 1998; Craft et al., 2005). Local supply of the LhG4 protein provided, multiple interspersed repeats of the operator elements (*pOp4; pOp6*) enables strong overexpression of a target gene in a cell type-specific manner (Craft et al., 2005).

Our work builds on these seminal studies by creating 19 well-characterized and stable driver lines targeting most cell types in *Arabidopsis* with a focus on the three main meristems of the plant, the root apical meristem (RAM), the shoot apical meristem (SAM), and the cambium. Of note, for several cell types such as the pith in the inflorescence stem or the xylem pole pericycle cells in the root, inducible expression systems were not available so far. The driver lines were generated employing the fast and flexible GreenGate cloning system (Lampropoulos et al., 2013), but are compatible with any vector/transgenic line in which the expression of an effector is under control of derivatives of the *pOp* promoter (Moore et al., 1998). An important feature of our driver lines is the presence of a fluorescent reporter amenable to live imaging, which allows monitoring the spatio-temporal dynamics of gene induction and may serve as a read-out for any effect on the respective tissue identity. Similarly, it allows to assess whether the expression of the effector has an impact on the transcriptional circuitries targeting the promoter it is expressed from. As trans-activation efficiently occurs in F1 plants derived from a cross between a driver and an effector line, the effect of a given expression cassette can be assessed relatively quickly in a wide range of cell types, demonstrating the usefulness of this resource for a broader research community. Moreover, testing the effect of genetic perturbations in a broad repertoire of individual tissues on a distinct developmental of physiological process seems feasible.

## Results

### Design of driver lines with cell type-specific expression of GR-LhG4

To generate a comprehensive set of driver lines expressing the chimeric GR-LhG4 transcription factor under control of cell type-specific promoters, we made use of the GreenGate cloning system, which enables quick modular assembly of large constructs (Lampropoulos et al., 2013). Our design included, on the same T-DNA, the coding sequence for an mTurquoise2 fluorescent reporter ((Goedhart et al., 2012) targeted to the endoplasmic reticulum (ER) through the translational fusion with an N-terminal signal peptide from sweet potato Sporamin A (SP, (Lampropoulos et al., 2013)) and the ER retention motif His-Asp-Glu-Leu (HDEL) under control of the pOp6 promoter (*pOp6:SP-mTurquoise2-HDEL*) (Figure 1). In our set up, the GR-LhG4 transcription factor is constitutively expressed dependent on the activity of a tissue-specific promoter (pTS). Consequently, GR-LhG4 activates the expression of the mTurquoise2 reporter and any other effector downstream of a *pOp* promoter after Dex treatment specifically in those tissues (Figure 1). We anticipate that the most utility can be obtained from this system if lines harbouring effector cassettes are crossed with driver lines and analyses are performed with F1 plants. However, other modes such as direct transformation of multiple driver lines or the introgression into different (mutant) backgrounds are also conceivable. Notably, even though the mTurquoise2 reporter is expressed from the same T-DNA as GR-LhG4, there is no mechanistic difference to the activation of an effector *in trans* (Figure 1).

**Figure 1.**
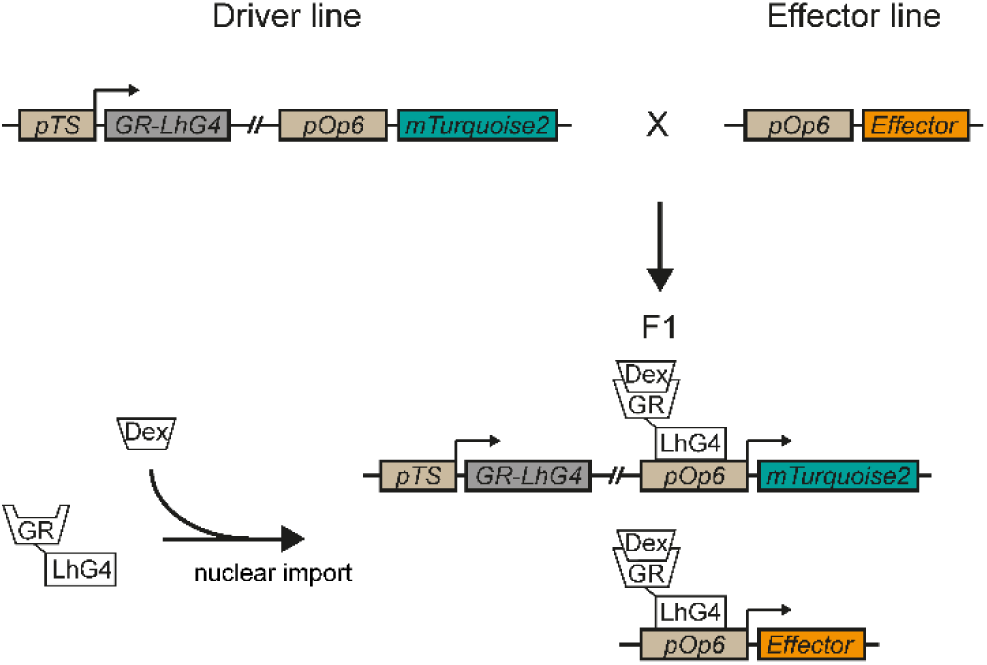
Overview of the Dex-inducible GR-LhG4/pOp system. In driver lines, expression of the synthetic transcription factor LhG4 is controlled by a tissue-specific promoter (*pTS*), whereas translational fusion with the ligand binding domain of rat glucocorticoid receptor (GR) prevents nuclear translocation in the absence of inducer (Dex). After crossing with an effector line harbouring a transcriptional cassette under control of a *pOp* promoter and addition of Dex, GR-LhG4 drives the expression of the effector as well as the mTurquoise2 reporter encoded by the driver line.

For establishing a rather comprehensive set of driver lines, we first selected respective tissue-specific promoters based on literature reports and our own expression data (Table 1). Subsequently, we generated stable transgenic driver lines in the *Arabidopsis thaliana* Col-0 background using 19 specific promoters that cover most cell types in the RAM, the SAM, and the cambium. Several of the promoters have been previously shown to work robustly in cell type-specific approaches (Marques-Bueno et al., 2016; Siligato et al., 2016). Next, we generated T3 lines in which the resistance to the selective agent sulfadiazine segregated as a single locus in the T2 generation or which showed a single insertion locus in SA-QPCR analyses (Huang et al., 2013).

**Table 1:** Overview of promoters utilized in this study.

| Promoter | Expression | Reference |
| --- | --- | --- |
| *pSCR* | Endodermis, quiescent centre in RAM, starch sheath in stem | (Di Laurenzio et al., 1996; Wysocka-Diller et al., 2000) |
| *pATHB-8* | Procambium, xylem precursors and columella in RAM | (Baima et al., 1995) |
| *pXPP* | Xylem pole pericycle cells | This study |
| *pAHP6* | Protoxylem precursors, pericycle, organ primordia in the SAM | (Mahonen et al., 2006; Besnard et al., 2014) |
| *pPXY* | (Pro-)cambium | (Fisher and Turner, 2007) |
| *pTMO5* | Xylem precursors | (Schlereth et al., 2010; De Rybel et al., 2013) |
| *pSMXL5* | Phloem (precursors) | (Wallner et al., 2017) |
| *pCASP1* | Endodermis | (Roppolo et al., 2011) |
| *pVND7* | Protoxylem (differentiating) in root, vessels in stem | (Kubo et al., 2005) |
| *pAPL* | Phloem (differentiating) | (Bonke et al., 2003) |
| *pNST3* | Fibres | (Mitsuda et al., 2007) |
| *pWOX4* | (Pro-)cambium | (Hirakawa et al., 2010) |
| *pLTP1* | Epidermis in stem | (Thoma et al., 1994) |
| *pAT2G3830* | Pith | (Valerio et al., 2004) |
| *pML1* | L1 layer, epidermis | (Lu et al., 1996) |
| *pCLV3* | SAM stem cells | (Fletcher et al., 1999) |
| *pREV* | SAM central zone | (Otsuga et al., 2001) |
| *pUFO* | SAM peripheral zone | (Levin and Meyerowitz, 1995) |
| *pCUC2* | Boundaries in SAM and leaf | (Aida et al., 1997) |

### Validation of the specificity of driver lines

To confirm the expected expression patterns in the root, driver lines were germinated on medium containing 30 μM Dex or DMSO, respectively, and analysed with Confocal Laser Scanning Microscopy (CLSM) five days after germination. In each case, we recorded mTurquoise2-derived fluorescence in longitudinal optical sections of the root meristem (Figure 2 and Supplemental Fig. 1) and, where appropriate, in cross sections through the meristem or the differentiation zone (Supplemental Fig. 2). To visualize expression in the shoot, lines were grown on soil in long day conditions and the aerial part of plants with 15 cm tall inflorescence stems were dipped either in tap water containing 10 μM Dex (Figure 3) or only the solvent DMSO (Supplemental Fig. 3). After 24 h, freehand sections of the stem were stained with propidium iodide (PI) to highlight xylem elements and analysed by confocal microscopy. To analyse expression in the SAM, inflorescence meristems of 15 cm tall plants were treated with Dex 48 hours before being dissected and imaged with CLSM, again using PI as a cell wall counterstain (Figure 4). Reporter gene activities were consistent with the expected patterns and strictly dependent on the presence of Dex (Supplemental Fig. 1, 3 and 4). In addition, the complete absence of reporter activity in tissues adjacent to cells in which activity was expected suggested that the chimeric GR-LhG4 protein does not move between cells.

**Figure 2.**
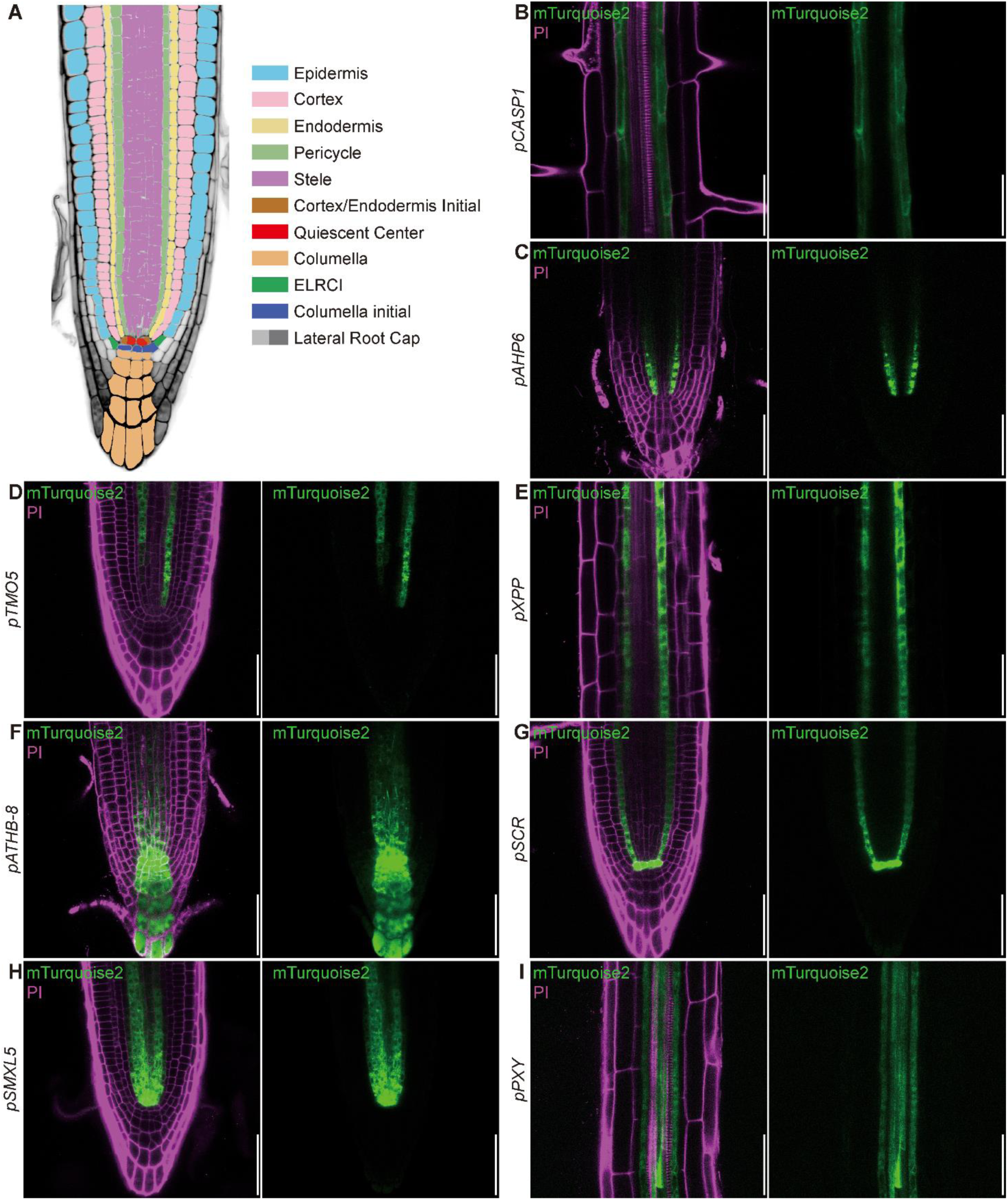
Analysis of induced driver lines in seedling roots. A, Schematic representation of root tissue layers. B-I, Induced driver line roots displaying fluorescence from propidium iodide-stained cell walls and the mTurquoise2 reporter (see Fig. 1 and Table 1). The indicated promoters mediate expression in the differentiating endodermis (B, *pCASP1*), phloem precursor cells and adjacent pericycle cells (C, *pAHP6*), xylem precursor cells (D, *pTMO5*), xylem pole pericycle cells (E, pXPP), stele initials, cortex/endodermis initial (CEI) and columella initials (F, *pATHB-8*), endodermis, CEI and quiescent centre (G, pSCR), stele initials, phloem and procambial cells (H, *pSMXL5*), and procambial cells (I, pPXY), respectively. Propidium iodide (PI) fluorescence is false-coloured in magenta and mTurquoise2 fluorescence in green. Bars = 50 μm.

**Figure 3.**
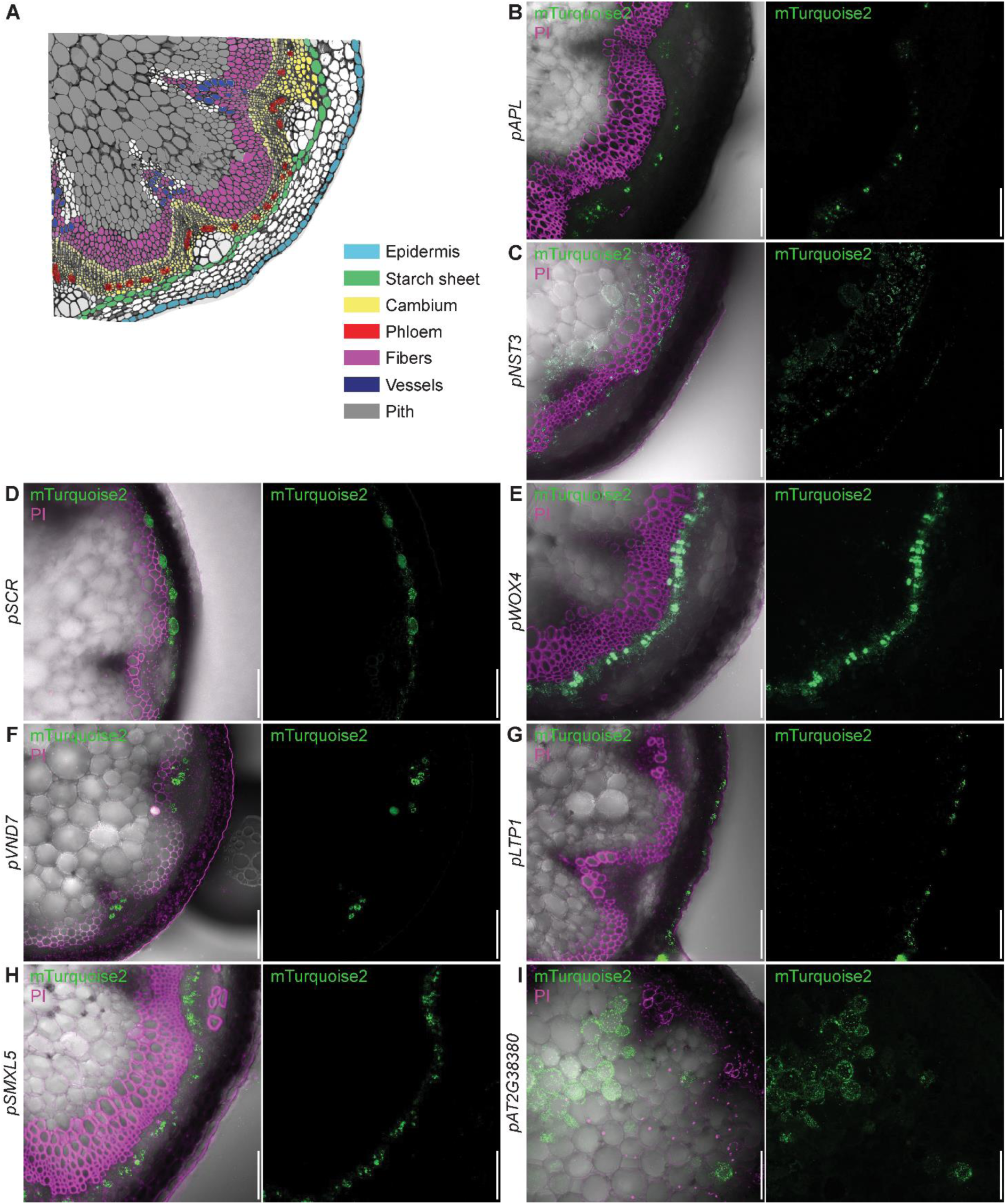
Analysis of induced driver lines in the stem. A, Schematic representation of inflorescence stem tissue layers. B-I, Induced driver line stems displaying fluorescence from propidium iodide-stained cell walls and the mTurquoise2 reporter (see Fig. 1 and table 1). The promoters mediate expression in differentiated phloem (B, *pAPL*), xylem fibres and interfascicular fibres (C, pNST3) starch sheath (D, pSCR), cambium (E, *pWOX4*), xylem vessels (F, *pVND7*), epidermal cells (G, *pLTP1*), the incipient phloem (H, *pSMXL5*), and pith (I, *pAT2G38380*), respectively. Propidium iodide (PI) fluorescence is false-coloured in magenta and mTurquoise2 fluorescence in green. Bars = 50 μm.

**Figure 4.**
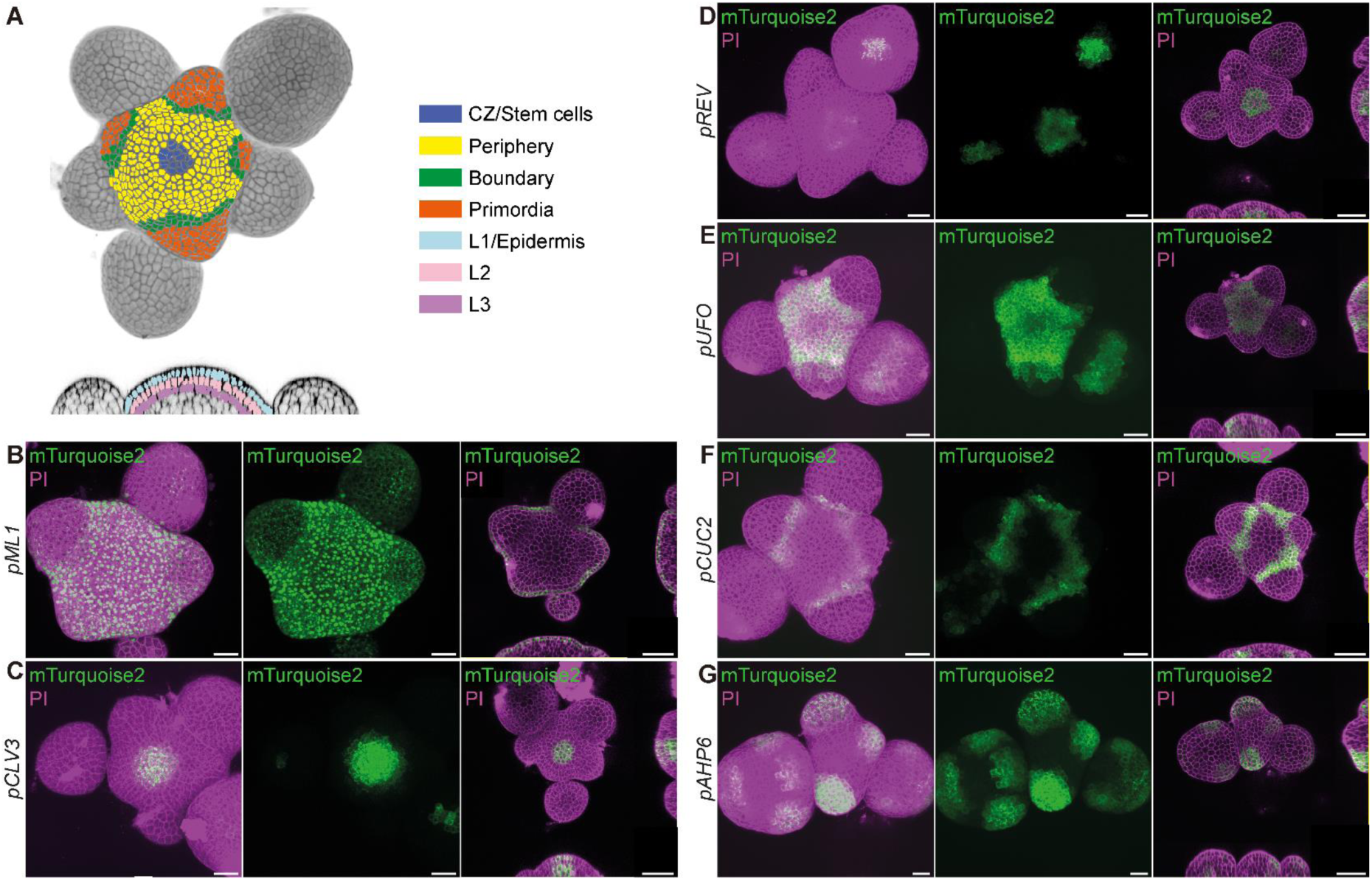
Analysis of induced driver lines in the shoot apical meristem (SAM). A, Schematic representation of cell identity domains in the SAM. B-G, Induced driver line stems displaying fluorescence from propidium iodide-stained cell walls and the mTurquoise2 reporter (see Fig. 1 and table 1). The indicated promoters mediate expression in the L1 layer/epidermis (B, *pML1*), the stem cell domain (C, *pCLV3*), the central zone (D, *pREV*), the peripheral zone (E, *pUFO*), the boundary domain (F, *pCUC2*), and organ primordia (G, *pAHP6*), respectively. Propidium iodide (PI) fluorescence is false-coloured in magenta and mTurquoise2 fluorescecne in green. Bars= 20 μm.

### Characterization of gene activation

We next tested whether dose-response and induction dynamics previously observed with the GR-LhG4 system (Craft et al., 2005) were recapitulated in our set up. To this end we germinated the *pSCARECROW* (*pSCR*) driver line mediating GR-LhG4 expression in the quiescent centre (QC) and the endodermis (Di Laurenzio et al., 1996; Wysocka-Diller et al., 2000) on plates containing solvent, 0.1 μM, 1 μM, 10 μM, and 100 μM Dex. Visualizing reporter fluorescence 5 days after germination indeed revealed increasing reporter activity with increasing Dex concentrations (Figure 5A), arguing for the possibility to fine tune gene expression by adjusting the levels of the inducer. We noticed that QC cells showed markedly stronger fluorescence compared to the endodermis, putatively reflecting higher promoter activity or GR-LhG4/reporter stability in the QC. We therefore quantified fluorescence separately in the QC cells and the endodermal initials (Figure 5C). Whereas the QC did not show significant difference in fluorescence intensity between any of the treatments, the endodermis responded in a linear fashion to increasing concentration of the inducer until saturation was reached between 10 μM and 100 μM of Dex (Figure 5C). Consequently, we concluded that, to fine tune gene expression by applying different Dex concentrations, the appropriate concentration range has to be determined for each promoter and cell type individually.

**Figure 5.**
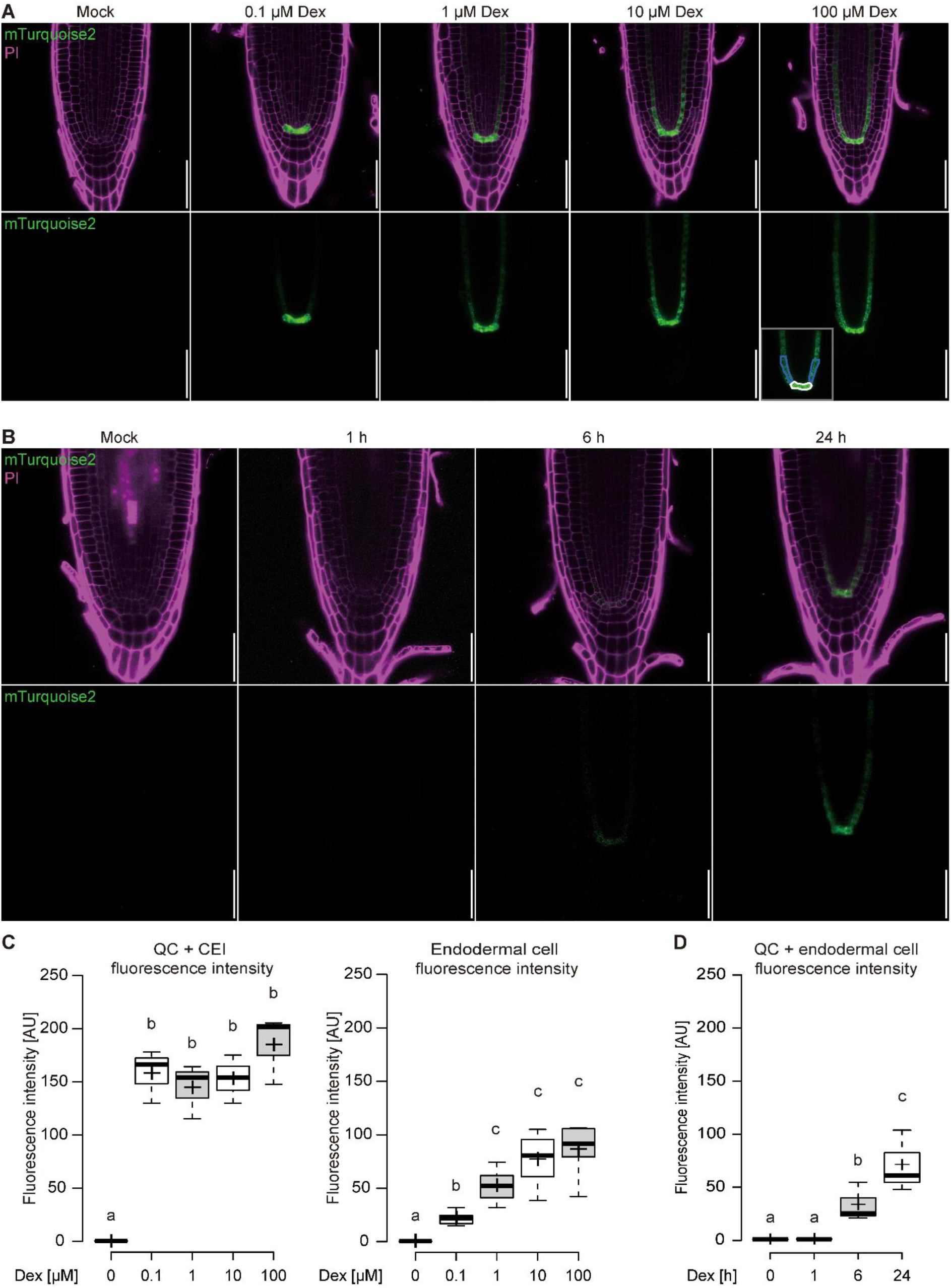
Dose-response and time course analysis of driver line seedling roots. A, The *pSCR* driver line was grown on 0, 0.1, 1, 10 and 100 μM Dex and imaged five days after germination. B, Time-course of pSCR driver line induction for 1, 6 and 24 hours with 10 μM Dex. C, Quantification of mTurquoise2 fluorescence intensity dose-response in quiescent centre cells and CEI (cells outlined in white in panel A). D, Quantification of mTurquoise2 fluorescence intensity of the first 3 endodermal cells after the CEI (cells outlined in blue in panel A). D, quantification of induction time-course (B) in quiescent centre cells, CEI and first 3 endodermal cells. Significance of difference is based on results of a two-tailed t test with p < 0.05, p < 0.01, p < 0.001, n=3 roots each. Bars = 50 μm.

To further assess induction kinetics, the *pSCR* driver line was germinated on plates with control medium and transferred onto plates containing 50 μM Dex after five days. As expected, a time-dependent increase of reporter activity was observed over a period of 24 hours (Figure 5B). Combined quantification of fluorescence in the QC and the endodermis initials detected reporter activity six hours after induction and activity values close to values of constitutive Dex treatment after 24 hours (Figure 5D). These observations suggested that six hours are sufficient to allow nuclear import of GR-LhG4, the induction of gene transcription, and initial protein translation, and that within 24 hours, protein levels reached a steady-state level.

To estimate the level of transcription mediated by the GR-LhG4/pOp system we employed a line expressing *PECTIN METHYLESTERASE INHIBITOR5* (*PMEI5*) (Wolf et al., 2012) under control of the strong and nearly-ubiquitous *35S* promoter (*p35S:PMEI5*). When comparing roots from the *p35S:PMEI5* line with roots from a Dex-treated GR-LhG4/pOp line conferring expression of the same *PMEI5* coding sequence in xylem pole pericycle cells (designated as *pXPP>GR>PMEI5* (Craft et al., 2005)), we observed *PMEI5* transcript levels similar to or slightly exceeding those in the *p35S:PMEI5* line (Supplemental Fig. 4). This was despite the fact that the *XPP* expression domain contains only ~six cell files in the root (Supplemental Fig. 2). Thus, we concluded that, although activating transcription in a very local manner, the GR-LhG4/pOp system can lead to strong expression in the respective cell types.

The ER-localized mTurquoise2 reporter present in our driver lines is transcribed from the same T-DNA that harbours the GR-LhG4 module (Fig. 1). To analyse the response of an independent T-DNA insertion carrying the *pOp6* element *in trans*, we generated a transgenic line carrying an ER-targeted mVenus reporter under control of the *pOp6* promoter (*pOp6:SP-mVenus-HDEL*) and crossed it with the *pSCR* driver line. The resulting F1 plants did not show any reporter activity when grown on plates without Dex (Fig. 6), again confirming that the GR-LhG4/pOp system is fully Dex-dependent. After Dex induction, we visualized both mTurquoise2 and mVenus fluorescence in the root and the stem and observed a complete congruence of both reporter activities (Fig. 6). Likewise, transgenic lines expressing a nucleus-targeted triple GFP fusion protein under the control of the *pOp6* promoter were generated and crossed with the *pCLAVATA3* (*CLV3*) driver line mediating expression in stem cells of the SAM (Fletcher et al., 1999). As expected, upon Dex induction, the 3xGFP-NLS signal was observed in a narrow domain at the tip of the SAM which also expressed the mTurquoise2 marker (Fig. 6). Together, these observations confirmed robust and specific trans-activation of transgenes in F1 plants.

**Figure 6.**
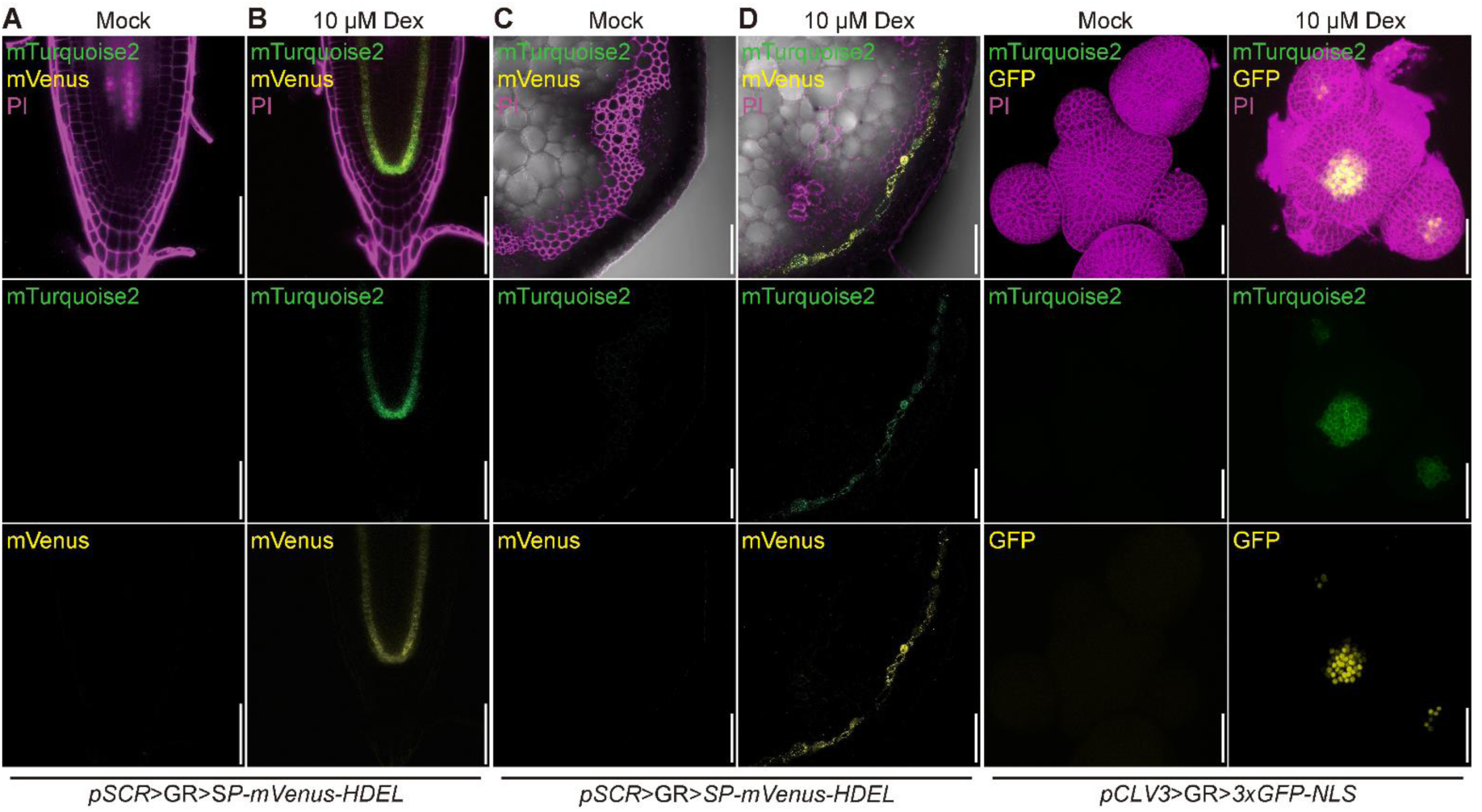
Induction of mTurquoise2 and mVenus/3xGFP fluorescence in root, stem and SAM of F1 plants from a driver line-effector line cross. Cells are counter-stained with PI (which, in the stem, highlights lignified vessel elements and fibres). Fluorescence channels are false-coloured. Bars = 50 μm for the root and the stem, 40 μm for the SAM.

### Cell-type specific induction of VND7 demonstrates efficacy of trans-activation

To explore the potential of our lines to mediate expression of a biologically active effector, we crossed the *pSCR* driver line with a line harbouring the VASCULAR RELATED NAC-DOMAIN PROTEIN7 (VND7) effector fused to the VP16 activation domain able to induce the formation of xylem vessels in a broad range of cell types (Kubo et al., 2005; Yamaguchi et al., 2010). F1 plants were grown on control medium for five days and then transferred to medium containing either 10 μM Dex or solvent. Five days later, fully differentiated vessel-like elements could be observed in the endodermis of both root and hypocotyl (Fig. 7), whereas in DMSO-treated controls xylem elements were clearly restricted to the stele. These results demonstrate that this resource for cell type-specific and inducible trans-activation can be used to study gene function with high spatio-temporal resolution.

**Figure 7.**
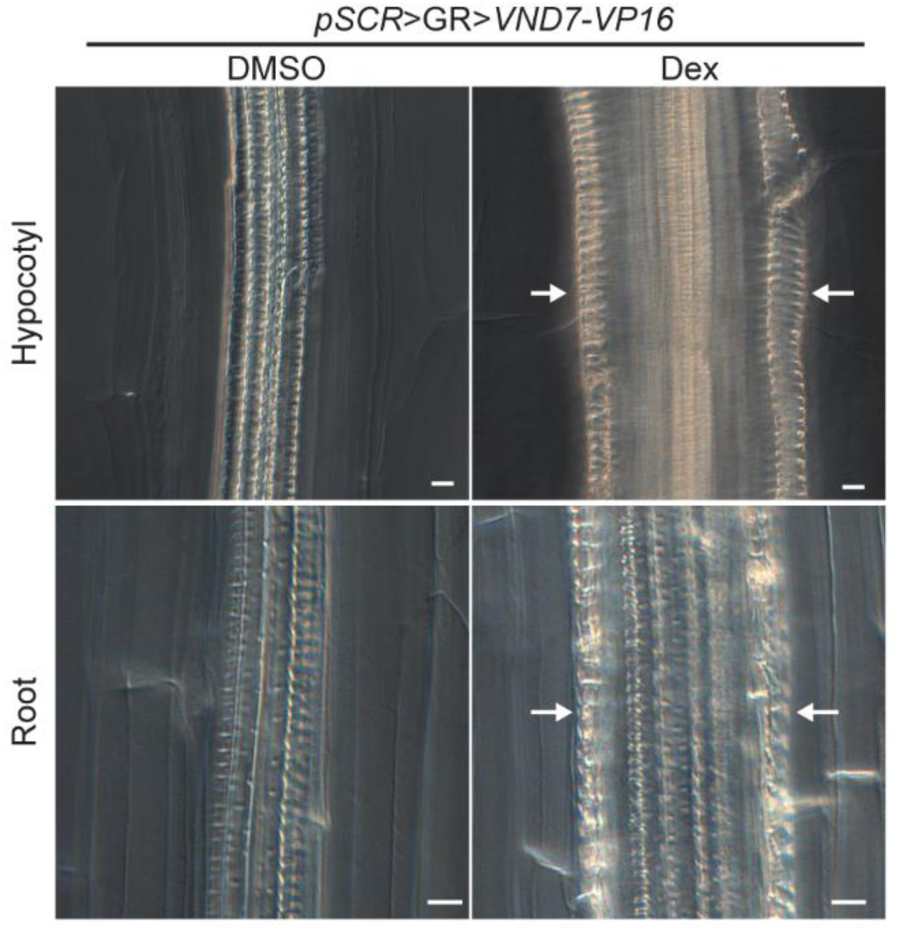
Cell-type specific induction demonstrates the efficacy of *trans-activation*. Plants expressing VND7-VP16 as an effector in the endodermal cells (*pSCR>GR>VND7-VP16*) show ectopic vessel formation (white arrows) after 5 days of Dex induction in both root and hypocotyl endodermis, in contrast to DMSO-treated plants. The spiral secondary cell wall thickening is observed after fixing and clearing the samples and visualized by DIC (differential interference contrast microscope). Bars = 20 μm.

## Discussion

In this study, we combined the proven efficacy of the well-established GR-LhG4/pOp expression system (Craft et al., 2005; Rutherford et al., 2005; Samalova et al., 2005) with the ease of cloning enabled by the GreenGate system (Lampropoulos et al., 2013) to provide a comprehensive toolbox for inducible, cell type-specific expression in Arabidopsis. The driver lines described here cover a large proportion of the known cell types in the three main meristems of the plant, the RAM, the SAM, and the cambium. Our analysis demonstrates that this system achieves non-leaky, adjustable, and robust trans-activation of effectors in the F1 generation after crossing with effector-carrying plants. Therefore, generating a line harbouring an effector cassette under control of the *pOp6* promoter should enable users to rapidly assess a battery of different expression regimes for a wide range of applications. In most cases, the effector might be the coding region of a gene one may want to miss-express in a spatially and temporally controlled manner, but other uses are conceivable, such as adjustable (pulsed) expression of reporters, domain specific knock-down through artificial microRNAs, cell-type specific complementation studies, the acquisition of cell type specific transcriptomes/translatomes/proteomes/epigenomes, or the local induction of genome editing, for example through expression of CRE recombinase or CRISPR/Cas9 modules (e.g. Birnbaum et al., 2003; Brady et al., 2007; Dinneny et al., 2008; Gifford et al., 2008; Mustroph et al., 2009; Deal and Henikoff, 2011; Hacham et al., 2011; Iyer-Pascuzzi et al., 2011; Petricka et al., 2012; Fridman et al., 2014; Adrian et al., 2015; Vragovic et al., 2015; Efroni et al., 2016; Kang et al., 2017). Thus, this system should be a valuable tool for the generation of inducible genetic perturbations to overcome the limitations of “endpoint” genetics and study genetic activities in specific tissue contexts.

### Design of the trans-activation system

Two-component trans-activation and chemically-induced gene expression systems have been widely used by plant biologist in the past. For example, a large collection of enhancer trapping lines based on the yeast Gal4 TF (Haseloff, 1999; Engineer et al., 2005) are an invaluable tool for constitutive, tissue-specific trans activation in Arabidopsis (Aoyama and Chua, 1997; Sabatini et al., 2003; Weijers et al., 2003; Swarup et al., 2005; Weijers et al., 2005). In addition, an inducible system based on Gal4/UAS has been devised (Aoyama and Chua, 1997) but appears to induce unspecific growth defects (Kang et al., 1999). Trans-activation based on LhG4 (Moore et al., 1998) shows only minimal detrimental effects on plant development, is thoroughly characterized and optimized (Moore et al., 1998; Baroux et al., 2005; Craft et al., 2005; Rutherford et al., 2005; Samalova et al., 2005; Moore et al., 2006) and has been used by the plant community in a number of studies (e.g. Schoof et al., 2000; Baroux et al., 2001; Eshed et al., 2001; Hay and Tsiantis, 2006; Nodine and Bartel, 2012; Sauret-Gueto et al., 2013; Hazak et al., 2014; Serrano-Mislata et al., 2015; Jiang and Berger, 2017). Parallel to the development of these tools for cell type-specific expression, a number of inducible systems have been conceived to enable temporal control of gene expression (Gatz et al., 1992; Weinmann et al., 1994; Caddick et al., 1998; Zuo et al., 2000). Subsequently, combining and optimizing the available technology has succeeded in generating tools to mediate inducible expression in a cell type-specific manner (Deveaux et al., 2003; Laufs et al., 2003; Maizel and Weigel, 2004; Craft et al., 2005).

For the generation of this resource, we build on ground-breaking previous work establishing the LhG4 system in combination with the GR ligand binding domain (Craft et al., 2005), which has since been proven to be an valuable resource (e.g. Reddy and Meyerowitz, 2005; Ongaro et al., 2008; Ongaro and Leyser, 2008; Heisler et al., 2010; Jiang et al., 2011; Dello Ioio et al., 2012; Merelo et al., 2016; Caggiano et al., 2017; Tao et al., 2017). For the generation of our driver lines, we exploited the power of the GreenGate cloning system (Lampropoulos et al., 2013). We were able to rapidly assemble a large number of constructs efficiently, circumventing the bottleneck previously imposed by the challenging generation of large DNA constructs with varying promoter elements, coding regions, and terminators. In our hands, the limiting factor in generating this resource was thus plant transformation, and obtaining single insertion, homozygous transgenic lines. As a general workflow, we aimed to generate at least 40 T1 transformants, then scored segregation rations of antibiotic/herbicide resistance in the T2 generation and maintained lines in which the resistance segregated as a single locus. These lines usually showed similar characteristics concerning the response to inducer and the expression levels achieved through trans-activation (based on fluorescence intensity). Nevertheless, reporter expression in any set of newly generated driver lines should be carefully assessed and compared with the literature and within lines, as genome integration in the vicinity of endogenous promoter and/or enhancer elements might influence the expression pattern. We occasionally observed widespread silencing in the T2 generation of the driver lines, which did not correlate with any particular module present in multiple constructs. In several cases, independent transformations of the same construct lead to stable expression, therefore the prevalence of silencing seemed to be influenced by the environmental conditions.

An important feature of our driver lines is the incorporation of a reporter amenable to live imaging, which can be used to monitor the induction and visualize the spatial expression domain. In addition, it allows to assess whether the expression of the effector has an impact on the transcriptional circuitries of the cell type it is expressed from. For some applications, the internal reporter of the driver lines might also serve as an inducible marker even in the absence of any further effector expression. We chose mTurquoise2 (Goedhart et al., 2012) as fluorescent reporter, since its spectral characteristics make it compatible with more widely used green and red fluorophores, and it displays high photostability, fast maturation, and high quantum yield. The fluorescent protein was N-terminally fused with a signal peptide and modified with a C-terminal HDEL motif to mediate retention in the ER, which is the preferable subcellular localization for a fluorescent reporter when cross sections through the highly differentiated cells of the stem are required.

### Trans-activation characteristics

Our system allows stringent temporal control of gene expression, as indicated by the lack of reporter expression in absence of the inducer Dex. Moreover, the trans-activated reporter faithfully reproduced previously described expression patterns associated with the respective 5’ regulatory regions, suggesting that the chimeric GR-LhG4 transcription factor is not cell-to-cell mobile. However, we noticed that in some cases trans-activation had the tendency to “flatten out” expression gradients observed with fusions of the same 5’ regulatory region with a reporter gene *in cis*. For example, expression driven from the *CLV3* promoter seemed broader than what was described in *pCLV3:XFP* lines, but consistent with a similarly designed pCLV3-driven trans-activation (Serrano-Mislata et al., 2015), possibly because the multiple binding sites of the *pOp6* promoter increase expression in cells where the *CLV3* promoter is only weakly active. Alternatively, high protein stability of the chimeric transcription factor, the reporter, or both, might cause prolonged activity of these proteins in cells that are already displaced from the stem cell region. This potential issue is less relevant for organs such as the root, where cells of one cell type also largely have the same clonal identity (Kidner et al., 2000; Costa, 2016).

Our experiments, in agreement with previous results, suggested that GR-LhG4/pOp-mediated trans-activation can achieve tissue-specific overexpression of the target gene, dependent on the concentration of inducer. However, the possibility of “squelching”, the sequestration of general transcription factors required for other processes by the LhG4 activation domain, must be taken into account at very high expression levels. Consistent with previous reports (Craft et al., 2005), our analysis of the *pSCR* driver line suggested that a linear dose-response over at least two orders of magnitude but the induction kinetics might be affected by genomic location of the transgene and thus should be empirically determined for each line. It should be noted that expression of effectors using LhG4/pOp systems can be quenched by adding Isopropyl β-D-1-thiogalactopyranoside (IPTG) (Craft et al., 2005), which would allow pulsing experiments. However, we did not test the effect of IPTG in our lines.

### Distribution of driver lines and DNA constructs

The lines described here, as well as DNA constructs, will be made available to the community through our website in a timely manner. Until completion of the website requests can be made by email. While GR-LhG4 and the sulfadiazine resistance gene are constitutively expressed, care should be taken to amplify seeds only from non-induced plants to minimize the chance of inducing post transcriptional gene silencing through high expression levels of the reporter (Schubert et al., 2004; Abranches et al., 2005).

## Material and methods

### Cloning

All constructs were produced by GreenGate cloning (Lampropoulos et al., 2013) using the modules described in Supplemental Table 1. The *E*co31l (*B*sal) sites of the *SCR, PXY* and *WOX4* promoters were removed by the QuickChange XL Site-Directed Mutagenesis Kit (Agilent Technologies, USA) using the primers in Supplementary Table 1 following the manufacturer instructions. The *E*co31I site of the *ATHB-8* promoter was removed by site-directed mutagenesis with primers containing the *E*co31I recognition site ‘GGTCTC’ and a four base overhang. The two fragments with the mutated *E*co31I recognition site were ligated afterwards.

The repetitive sequences of the *pOp* promoter increase the likelihood of recombination events while amplifying the plasmids. To discriminate against clones with shorter *pOp* sequences, we designed primers that bind in the short flanking sequences at the beginning and end of the *pOp6* (*pOp6*_F TGCATATGTCGAGCTCAAGAA; *pOp6*_R CTTATATAGAGGAAGGGTCTT) for PCR amplification and size assessment through gel electrophoresis. Final constructs were always confirmed by sequencing in *E. coli* and *Agrobacterium*. The occasional recombination events were only detected in *E. coli*.

### Plant material and growth conditions

All constructs were transformed by the floral dip method (Clough and Bent, 1998) as modified by (Zhang et al., 2006) into *Arabidopsis thaliana* Col-0. Transformed seeds were selected on ½ MS plates containing 1.875-3.75 μg/ml sulfadiazine. All plants were grown in long day conditions (L16:D8) at 22°C. For root analysis, plants were grown in 1% sucrose, 0.9% agar ½ MS plates. For the induction treatments in plates, the seeds were sown on plates containing dexamethasone (Sigma_D4903, St. Louis, Missouri, United States) in the indicated concentration while the same volume of DMSO (D139-1, Fisher Scientific, UK) was added for the mock control. For the trans-activation experiment, seeds were sown on plates without Dex and seedlings were transferred to Dex-containing plate at 1, 6 and 24 hours before imaging five days after germination. For analysis of the stem, the aerial parts of 15 cm tall plants were dipped for 30 s in either tap water containing 10 μM Dex with 0.02% Silwet L-77 (Kurt Obermeier GmbH & Co. KG, Bad Berleburg, Germany) or water with the same volume of DMSO with 0.02% Silwet. After 24 hours, free-hand sections of the stem were performed with a razor blade. Sections were transferred to a small petri dish (35/10mm, Greiner Bio-One GmbH, Germany) with 0.25 mg/ml of propidium iodide for 5 min and mounted in microscope slides to be visualized by CSLM. For SAM imaging, the inflorescence meristems of 25-30 DAG plants were sprayed with 10 μM Dexamethasone, whereas an equal volume of DMSO was added to the mock controls. 48 h after the treatment, the inflorescent meristems were dissected by cutting of the stem, flowers and buds. The SAM was stained in 0.25 mg/ml propidium iodide (Sigma-Aldrich, P4170) for 5 min and mounted in a 3 % agarose small petri dish (35/10mm, Greiner Bio-One GmbH, Germany) and visualized by CLSM.

### Microscopy

Root samples have been imaged by using Leica TCS SP5 laser scanning confocal microscope with a HCX PL APO lambda blue 63x water immersion objective. mTurquoise2 fluorophor was excited by argon laser at 458 nm and emission was collected between 460 and 516 nm. mVenus fluorophor was excited by at 514 nm and emission was collected between 520 and 580 nm. Cells were counter-stained by propidium iodide (Sigma-Aldrich, P4170) and imaged with 488 nm for excitation and emission was collected between 590 and 660 nm.

For stem and SAM samples we used a Nikon (Minato, Tokyo, Japan) A1 Confocal with a CFI Apo LWD 25x water immersion objective. The PI counter-stained cells were imaged with 561 nm for excitation and 570-620 nm for emission. The mTurquoise2 fluorescence acquired using excitation at 405 nm and emission was collected between 425-475 nm. For the trans-activation experiments, the 3xGFP-NLS signal in the SAM was imaged with 488 nm for excitation and 500-550 nm for emission. In the root, the mVenus was excited with 514 nm and the emission was collected between 500-550 nm.

For visualization of the xylem, plants were germinated in ½ MS plates and 5 DAG were transferred to either 10 μM Dex or mock containing ½ MS plates. To visualise the ectopic xylem formation, 5 DAI plants were collected and fixed overnight in a 1:3 acetic acid:ethanol solution. Then, they were cleared in a 8:1:2 Chloral hydrate:glycerol:water solution for at least 3 hours. Samples were mounted in microscope slide containing 50 % glycerol solution and brightfield images were obtained using an Axioimager M1 microscope equipped with an AxioCamHRc (Carl Zeiss, Jena, Germany).

## Acknowledgements

The authors thank the members of the Greb, Wolf, and Lohmann labs, and Alexis Maizel for discussion and support. This work was supported by the German Research Foundation DFG (grant WO 1660/6-1 to S.W. and grant GR 2104/4-1 to T.G.) and an ERC Consolidator grant (PLANTSTEMS, 647148) to T.G. SW is supported by the DFG through the Emmy Noether Programme (WO 1660/2-1).

---

**Supplemental Figure 1.**
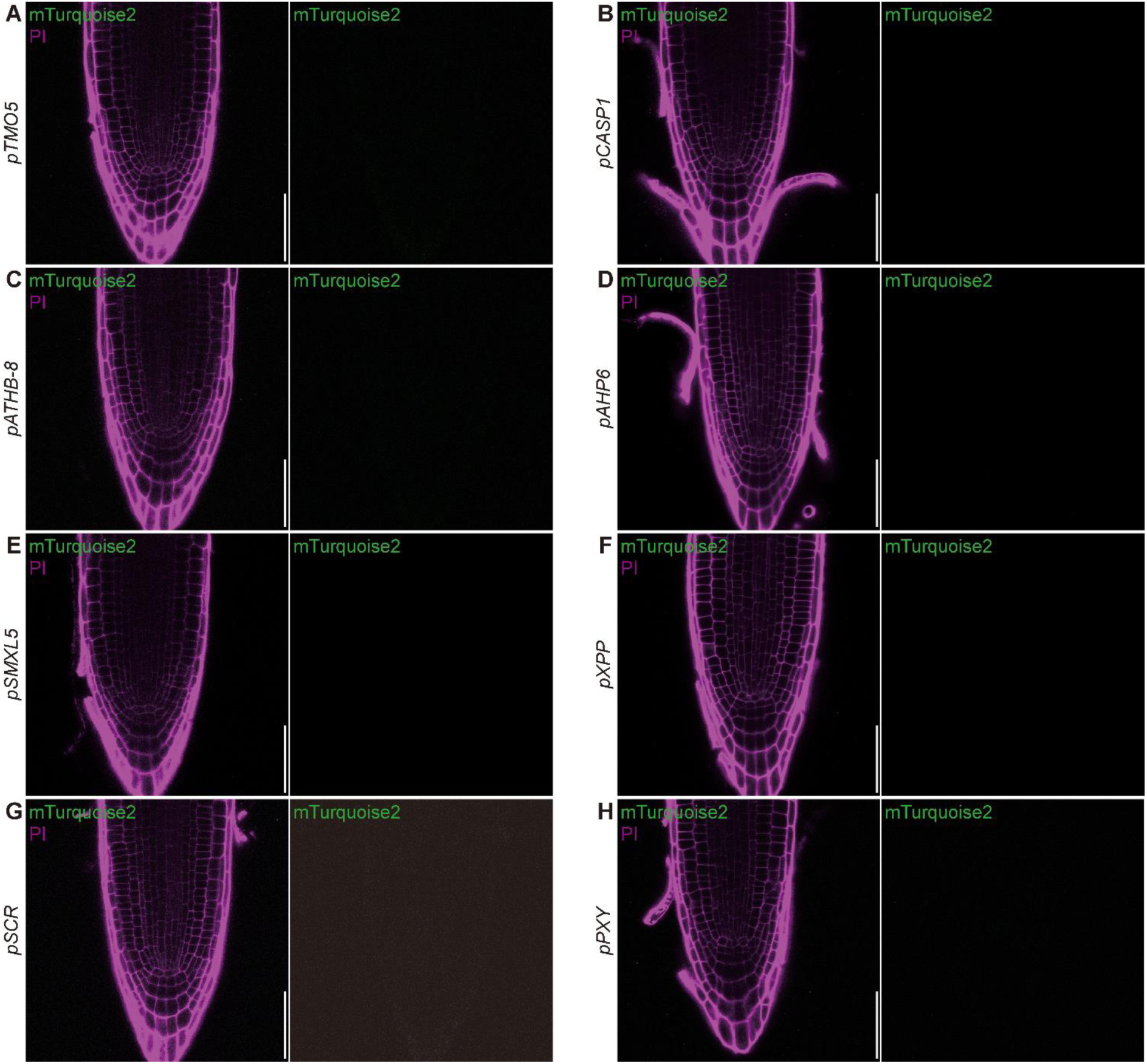
Analysis of DMSO-treated mock control for driver line seedling root induction 5 DAG. Cells are counter-stained with PI. Bars = 50 μM.

**Supplemental Figure 2.**
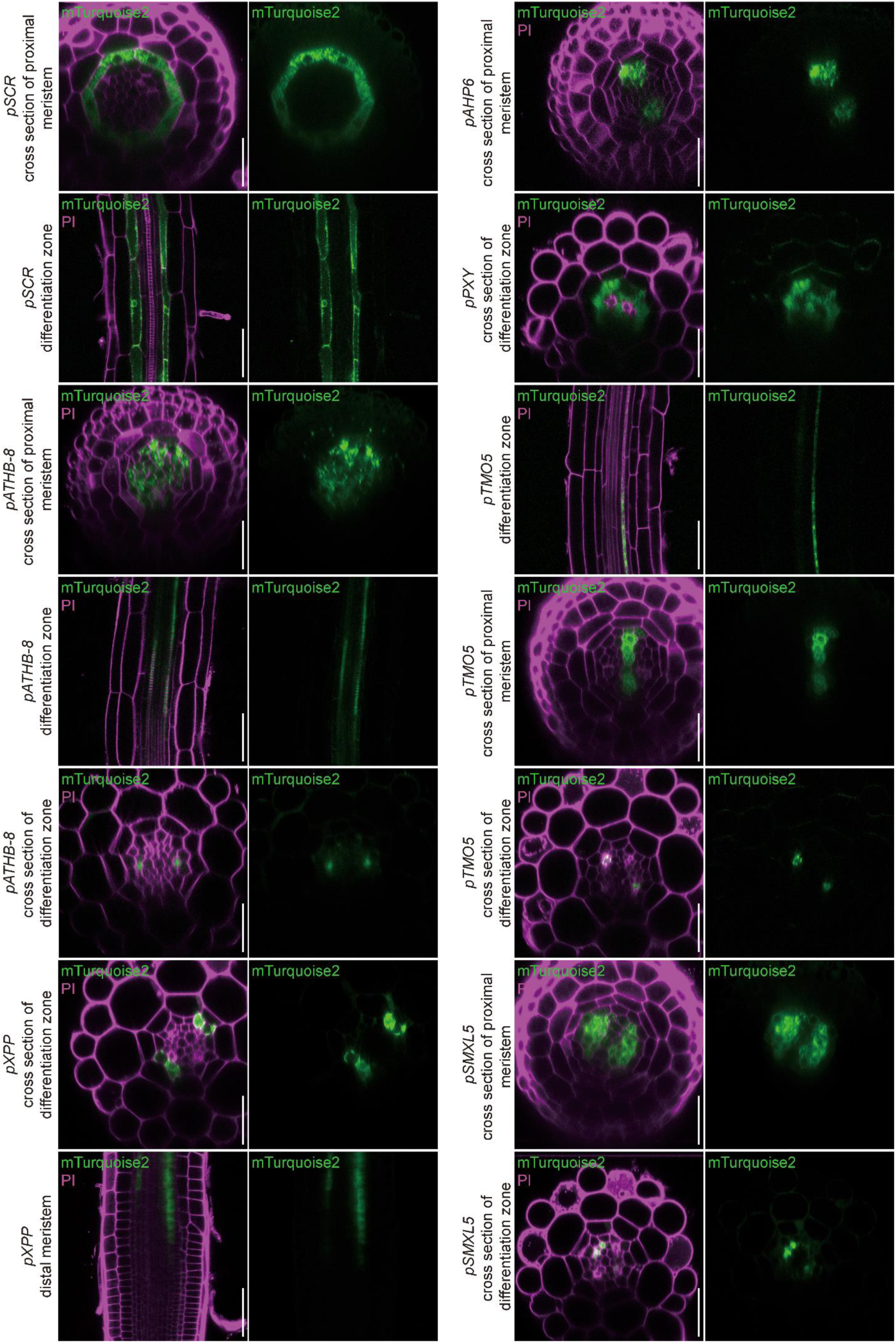
Analysis of induced driver lines in 5 DAG seedling root: cross section images and longitudinal images of proximal root apical meristem and differentiated regions showing signal in tissues at different developmental stages. Bars = 50 μM.

**Supplemental Figure 3.**
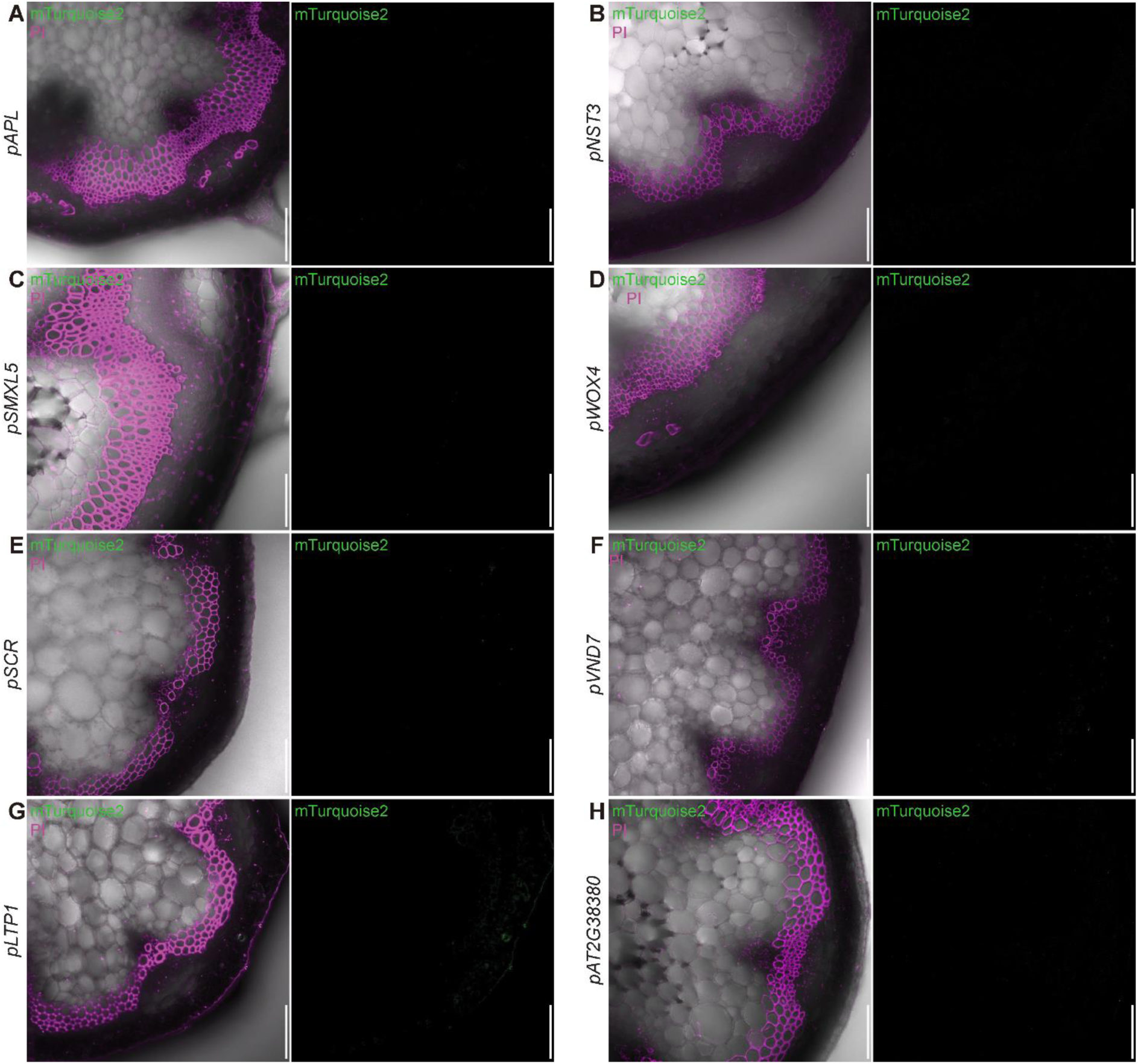
Analysis of DMSO-treated driver lines in the stem. A-H, In all driver lines mTurquoise2 fluorescence was absent in the mock treated samples. Cells are counter-stained with PI. Bars = 50 μM.

**Supplemental Figure 4.**
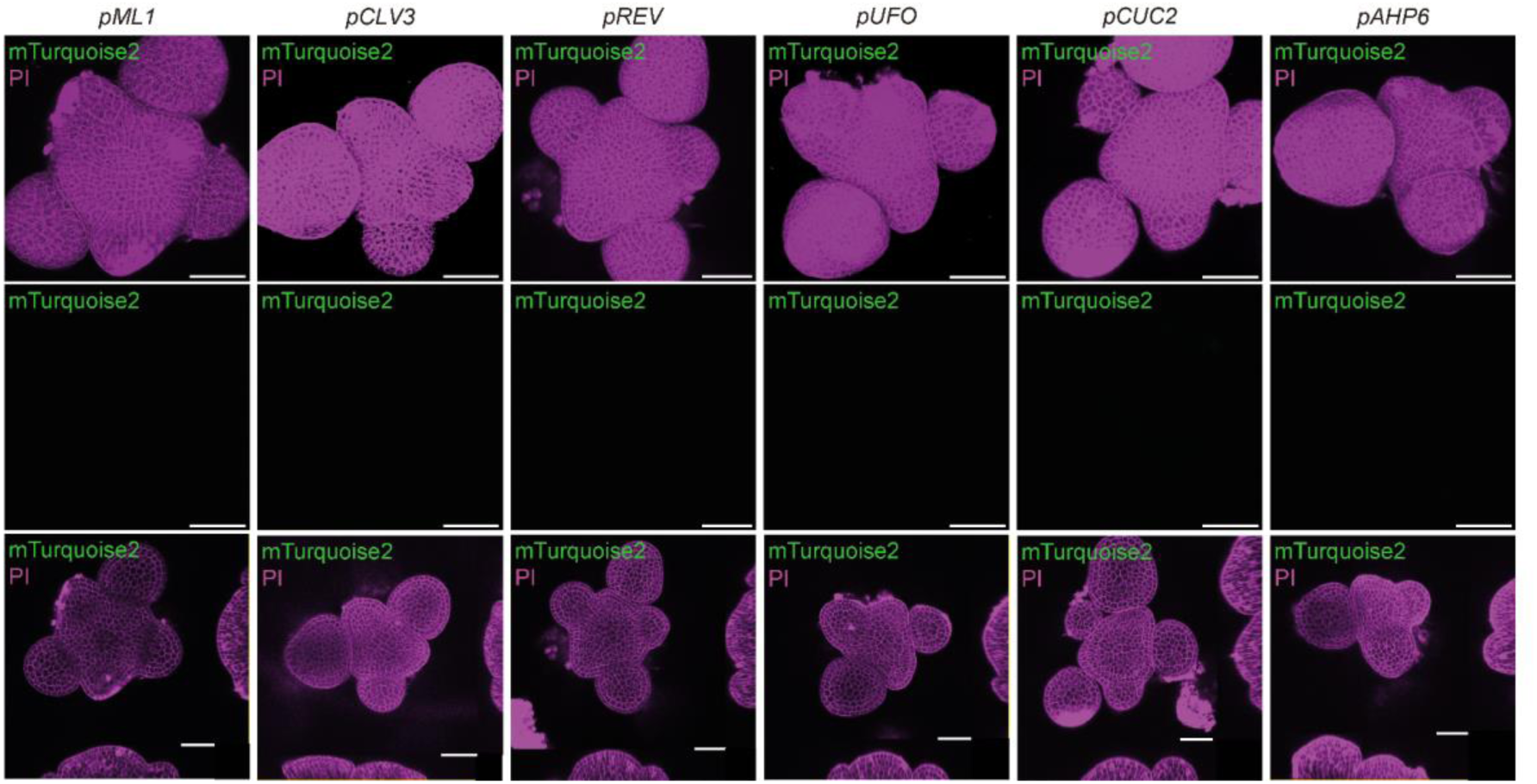
Analysis of DMSO-treated driver lines in the SAM. A-F, In all driver lines mTurquoise2 fluorescence is absent after DMSO-mock treatment for 48 hours. Cells were counter-stained with PI. Bars = 40 μM.

**Supplemental Figure 5.**
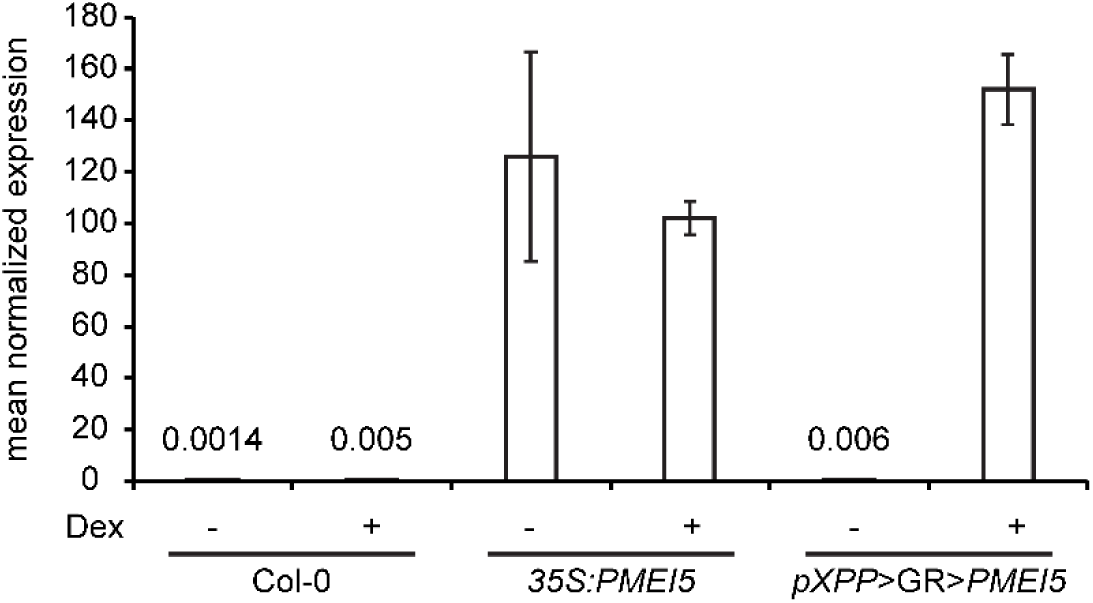
Qunatification of GR-LhG4-mediated trans-activation. qRT-PCR analysis of *PMEI5* expression driven by *pXPP* in xylem pole pericycle cells after Dex induction showed comparable expression levels to a line expressing PMEI5 under control of the ubiquitous *35S* promoter (PMEIox, (Wolf et al., 2012).

